# From time-course expression to gene regulation: direct linear ODE inference without finite-difference approximation

**DOI:** 10.64898/2026.05.18.726023

**Authors:** Xiaoqing Huang, Andersen Ang, Aatman Pushkarkumar, Kun Huang, Teresa M. Przytycka, Yijie Wang

**Affiliations:** Department of Biostatistics and Health Data Sciences, IUSM, Indianapolis, IN 46202, USA; School of Electronics & Computer Science, University of Southampton, SO17 1BJ, UK; Computer Science Department, Indiana University Bloomington, Bloomington, IN 47408, USA; Intramural Research Program, National Library of Medicine, NIH, Bethesda, MD, 20894, USA

## Abstract

Inferring gene regulation from time-course expression profiles is essential for understanding how cells transition between states during development, differentiation, and disease progression. Existing approaches often model expression dynamics with ordinary differential equations (ODEs). However, due to the computational complexity of directly solving these ODE models, most methods rely on finite-difference approximations of temporal derivatives, which can amplify measurement noise, introduce discretization bias, and lead to unstable or biased parameter estimates.

To fill this gap, we develop the first computational method to directly learn a linear ODE model for gene regulation inference without relying on finite-difference approximations. We first formulate an optimization problem that directly exploits the closed-form solution of the linear ODE system. We then solve this problem via gradient descent, deriving analytical gradients with respect to the model parameters; these gradients involve matrix exponentials and integrals, which are challenging to directly compute. To make the computation efficient, we further use high-order Taylor approximations of the gradients whose truncation error is on the order of machine precision. In addition, we establish theoretical results demonstrating an inherent, non-vanishing gap between our exact solution and solutions derived from finite-difference approximations, which underscores the theoretical advantages of our approach. Finally, we demonstrate that our method consistently outperforms competing approaches on both simulated data and real-world scRNA-seq datasets in terms of AUROC. Our source codes can be accessed here: https://github.com/EJIUB/ExactLinearODE

## 1 Introduction

Gene regulation inference is a longstanding challenge in systems biology. Many methods (reviewed in [1, 2]) infer regulatory relationships under a steady-state assumption, meaning that gene expression does not change appreciably over the measurement window. These steady-state gene regulation inference methods typically leverage bulk or single-cell expression profiles acquired at a single time point—rather than temporal measurements—to infer equilibrium regulatory structure. Recent advances in biotechnology enable time-course measurements of bulk expression and single-cell mRNA profiles along developmental or differentiation trajectories, making it possible to move beyond steady-state assumptions and study gene regulation as a dynamic process.

In this paper, we study the dynamic gene regulation inference methods for time-course expression data. The time-course expression data may arise from bulk mRNA profiles sampled at multiple time points, or from single-cell measurements ordered along developmental or differentiation trajectories via pseudotime. A large body of work approaches this problem with model-based frameworks—most notably Granger-causality models [3, 4, 5, 6, 7] and quantitative ordinary differential equation (ODE) formulations [8, 9, 10, 11, 12, 13]—that aim to capture the temporal structure of gene expression. In these frameworks, the regulatory network is inferred by optimizing model parameters against time- or pseudotime-resolved expression.

Granger-causality models [3, 4, 5, 6, 7] employ a range of computational formulations (linear vcector autoregressive model [3], kernel method [7], and deep learning models [6], etc) to test whether incorporating the past expression of candidate regulators improves the prediction of a target gene’s future expression beyond what is explained by the gene’s own expression history. When incorporating a transcription factor’s (TF) past expression significantly improves the prediction of a target gene’s future expression, the TF is considered a putative regulator of the target. This framework is appealing because it is conceptually straightforward for modeling multivariate temporal dependencies. However, Granger-causality models do not model the underlying dynamics that govern how expression evolves over time.

ODE-based approaches [8, 9, 10, 11, 12, 13], in contrast, explicitly parameterize the underlying dynamical law from time-course expression data. However, due to computational complexity, nearly all of them rely on finite-difference approximation to estimate the rate of change in the ODE functions. This is problematic because finite-difference approximations amplify measurement noise, introduce discretization bias when the sampling interval *Δt* is not small, and perform poorly with irregular, sparse or stiff dynamics, ultimately leading to unreliable derivatives and unstable, biased ODE parameter estimates.

To fill this gap, we develop the first computational method to learn a linear ODE system for gene regulation inference without relying on finite-difference approximations. Specifically, we model gene regulation and expression dynamics using a linear ODE system with a steady state, and exploit its closed-form solution via the matrix exponential to formulate an optimization problem for estimating the ODE parameters. We then solve this optimization problem using gradient descent. A key challenge is that the analytical gradients of the ODE parameters involve matrix exponentials and integrals, making them difficult to compute directly. To address this, we propose to use the high-order Taylor approximation of the gradients and show that this approximation is extremely accurate, with error on the order of machine precision. In addition, we provide theoretical results showing that there is an inherent, non-vanishing gap between our exact solution and those from finite-difference approximations, thereby demonstrating the theoretical superiority of our method. Finally, we demonstrate that our method consistently outperforms existing methods on both simulation studies and three time-course scRNA-seq datasets in terms of Area Under the Receiver Operating Characteristic (AUROC) scores.

## 2 Related works and our contributions

Gene-regulation inference from time-course expression data has been extensively studied. Existing approaches fall into two broad categories: causality-based methods and ODE-based methods. Causality-based methods [3, 4, 5, 6, 7] are grounded in the Granger-causality framework. They test whether past values of one TF improve the prediction of another gene’s future expression. For example, SWING [14] uses multivariate Granger causality with sliding-window regression; BETS [4] infers Granger causality via bootstrap elastic-net regression; CGC-2SPR [5] applies a two-step prior with ridge regularization; SINGE [7] employs kernel-based Granger causality; and GRANGER [6] employs deep-learning models to estimate Granger causality.

ODE-based methods, in contrast, model expression dynamics by specifying the rate function *f* (·) in 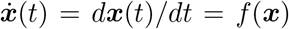, where ***x***(*t*) is the expression of genes at time *t*. Linear-function approaches assume *f* (***x***) = *A****x***. Examples include SCODE [8] and GRISILI [9]; PROB [11] estimates the linear coefficients via Bayesian Lasso; SCOUP [15] uses an Ornstein–Uhlenbeck process; and TSNI [16] uses a discrete linear ODE formulation. Nonlinear-function approaches allow *f* (***x***) to be nonlinear, e.g., random-forest regressors in dynGENIE3 [10], Hill-type kinetics in Ocone’s method [13], and Gaussian-process models in BINGO [12]. Most of the above methods do not directly solve the continuous ODE due to computational cost; instead, they approximate derivatives 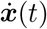with finite-difference approximation

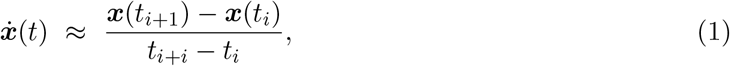

where ***x***(*t*_*i*_) and ***x***(*t*_*i*+1_) are the expression measurements at adjacent times *t*_*i*_ and *t*_*i*+1_.

For completeness, we also note that several Neural-ODE-based methods [17, 18] have been developed; however, they aim to model the dynamics of gene regulation rather than inference.

### Contribution

Our contributions are in two folds:

I. We solve a linear ODE model via solving an optimization problem formulated using its exact analytic closed-form solution rather than the finite-difference approximation 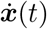 in Eq. (1) (details in section 3).
II. We establish theoretical results showing that there is an inherent, non-vanishing bias between our exact method and methods using the finite-difference approximation (details in section 4).

## 3 Methodology

In this section, we model gene expression dynamics using a linear ODE system and learn its parameters via optimization. We first introduce our linear ODE model in section 3.1 and formulate an optimization problem to learn its model parameters in section 3.2. We then derive the analytical closed-form gradients (involving matrix exponential and integration) for the optimization in section 3.3 and their Taylor approximation in section 3.4. Finally, we describe the gradient descent method to find the optimal model parameters in section 3.5.

### 3.1 Modeling gene regulation using a linear ODE system with steady states

In this work, we focus on inferring regulatory interactions among transcription factors (TFs), as done in [8, 9]. Let ***x***(*t*) ∈ ℝ^*n*^ denote the vector of *n* TFs’ expression levels at time *t* (as shown in Fig. 1(a)). We model their expression dynamics by a linear ODE system around a steady state ***x***_*_ (as shown in Fig. 1(b)):

**Fig. 1:**
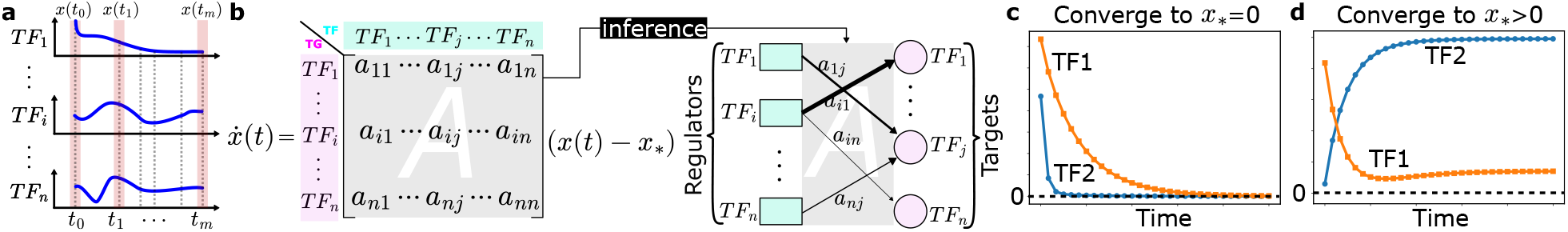
Overview of our linear ODE system. (a) Time-course expression data with irregular intervals. (b) Linear ODE system with steady states and how we use |*a*_*ij*_| in *A* to infer the regultory interactions. Edge thickness corresponds to the magnitude of |*a*_*ij*_|, which encodes confidence (thicker edges indicate higher confidence). (c) An example of TFs’s expression in a linear ODE system converge to ***x***_*_ = 0. (d) An example similiar to (c) but ***x***_*_ *>* 0. TFs’ expression converge to positive values.

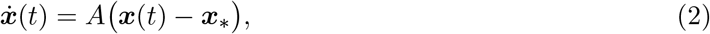

where ***x***_*_ ∈ ℝ^*n*^ and *A* ∈ ℝ^*n×n*^ are the model parameters that need to be learned from time-course expression measurements. Specifically, ***x***_*_ is the equilibrium (steady-state) expression to which the *n* TFs’ expressions converge, and *A* is the interaction matrix, where *a*_*ij*_ quantifies how the expression of TF *j* influences the rate of expression change of target TF *i* (as shown in Fig. 1(b)). Given ***x***_*_, *A*, and ***x***(0), we can compute ***x***(*t*) for any *t* by the closed-form solution of the linear ODE system (2)

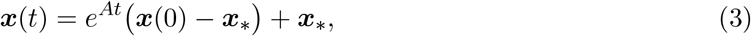

where the matrix exponential

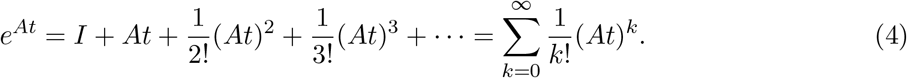

A key distinction of our model (2) from existing linear ODE models such as SCODE and GRISILI is that we explicitly consider ***x***_*_ in the model rather than setting ***x***_*_ = **0**. As illustrated in Fig. 1(c), imposing ***x***_*_ = **0** forces all TF trajectories to decay toward zeros, whereas explicitly modeling ***x***_*_ allows trajectories to converge to non-negative equilibrium levels (Fig. 1(d)). Clearly, considering ***x***_*_ *>* **0** avoids the biologically implausible implication that all TFs converge to zero, and is expected to yield better fitting and interpretability.

### Infer regultory interactions from *A*

The model parameter *A* can be used to infer regulatory interactions (as shown in Fig. 1(b)). We interpret *A*’s entries |*a*_*ij*_| as regulatory strengths and rank candidate regulatory interactions by |*a*_*ij*_|, where a larger magnitude indicates greater confidence in the presence of the corresponding interaction.

### 3.2 Learning the linear ODE system via optimization from expression dynamics

Given a time-course TF expression profile {(***x***_*i*_, *t*_*i*_) ∈ ℝ ^*n*^ *×*ℝ : *i* = 0, …, *m*}, where ***x***_*i*_ ∈ ℝ ^*n*^ is a vector that denotes the observed expression of *n* TFs at (pseudo)time *t*_*i*_. We can learn model parameters ***x***_*_ and *A* in (2) by solving the following optimization problem:

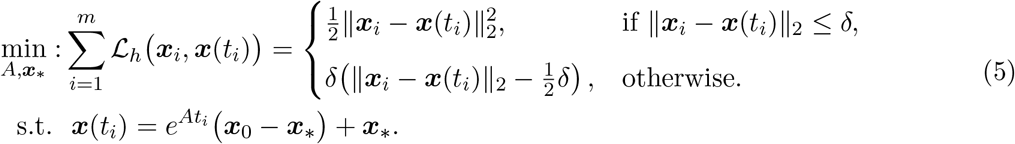

The above optimization (5) aims to find the optimal ***x***_*_ and *A* to minimize the Huber loss ℒ _*h*_ between observed expression ***x***_*i*_ and the corresponding model prediction ***x***(*t*_*i*_). We use the Huber loss *L*_*h*_ since it is less sensitive to outliers (where ||***x***_*i*_ − ***x***(*t*_*i*_) ||_2_ *> δ*), which is beneficial when fitting real-world noisy expression data, such as scRNA-seq data subject to dropout.

Directly solving optimization (5) is challenging; therefore, previous works [8, 9, 10, 11] use finite difference approximations (1) to estimate. In this paper, we directly solve it using the gradient-based method, where the main difficulty lies in computing 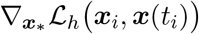 and ∇_*A*_ℒ _*h*_ (*x*_*i*_, *x*(*t*_*i*_)), as detailed in the next section.

### 3.3 Derivation of the closed-from gradient using Fréchet directional derivative

We now derive the gradients 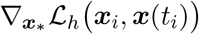 and ∇_*A*_ℒ _*h*_ (***x***_*i*_, ***x***(*t*_*i*_)). Let us first rewrite the loss function as

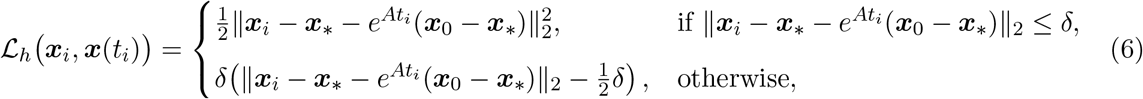

which is derived by substitulting ***x***(*t*_*i*_) in the loss function ℒ _*h*_ in (5) with 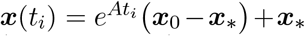.

Let us first derive the gradient 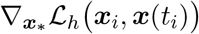. Let 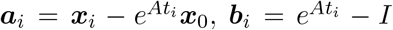, then 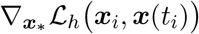 is

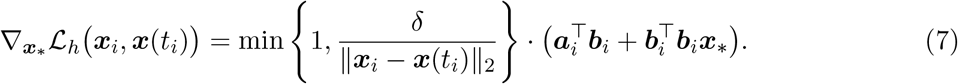

The derivation of 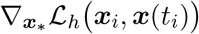 is straightforward and is therefore left to the audience.

We next derive the gradient ∇_*A*_ℒ_*h*_ (***x***_*i*_, ***x***(*t*_*i*_)), which is challenging because it involves the derivative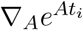 of the matrix exponential. To handle this, we use the Fréchet directional derivative of the matrix exponential [19], yielding the expression for ∇_*A*_ℒ*h(****x***_*i*_, ***x***(*t*_*i*_))stated in lemma 1.

#### Lemma 1

*Let* ***c***_*i*_ = ***x***_*i*_ − ***x***_*_, ***d*** = ***x***_0_ − ***x***_*_, *and* 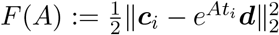, *then ∇*_*A*_ℒ_*h*_ (***x***_*i*_, ***x***(*t*_*i*_)) *is*

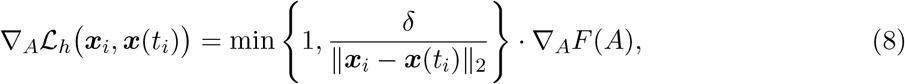

*where* 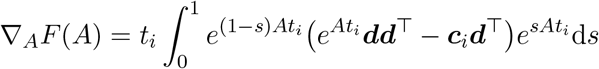

The proof of Lemma 1 is provided in the supplementary materials.

### 3.4 Gradient estimation using Taylor approximation for infinite series expansions

Even though we have closed-form expressions for 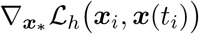 and ∇_*A*_ℒ_*h*_ (***x***_*i*_, ***x***(*t*_*i*_)) in (7) and (8), respectively, efficiently computing these gradients in practice is still challenging.

#### Computation of 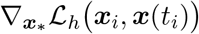

Let us compute 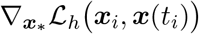 in Eq. (7), where we need to compute matrix exponential 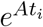 in both ***a***_*i*_ and ***b***_*i*_. The matrix exponential is defined in Eq. (4) as an infinite sum, which is impossible to implement; thus it is usually computed numerically by approximation with sufficiently small error. A large effort in numerical analysis has been dedicated to designing efficient computation of matrix exponential [20, 19, 21, 22]. We consider a degree-18 Taylor approximation of the matrix exponential *e*^*A*^ as

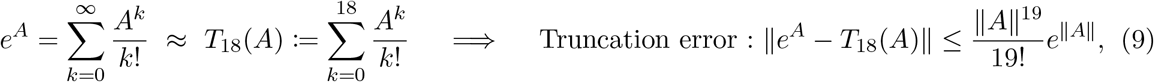

which the Taylor approximation assumes the matrix 2-norm *A* _2_ is small^5^. The degree-18 Taylor approximation is extremely accurate (see Theorem 11.2.4 in [21]), with an error on the order of machine precision (error estimate and implementation details in the supplementary materials).

#### Computation of ∇_*A*_ℒ_*h*_ (*x*_*i*_, *x*(*t*_*i*_))

The main computational task in evaluating ∇_*A*_ℒ_*h*_ (***x***_*i*_, ***x***(*t*_*i*_)) is to compute ∇_*A*_*F* (*A*) as defined in Lemma 1. In Lemma 1, the analytical closed form of ∇_*A*_*F* (*A*) is difficult to work with because it involves both a matrix exponential and an integral. Following the approach of Al–Mohy and Higham [19], we instead derive ∇_*A*_*F* (*A*) from its Fréchet derivative *L*_*F*_ (*A, E*), which admits a series expansion representation.

We consider 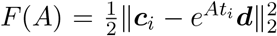 and compute the Fréchet directional derivative *L*_*F*_ (*A, E*) of *F* at *A* in the direction *E*. Let 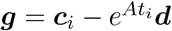. Then

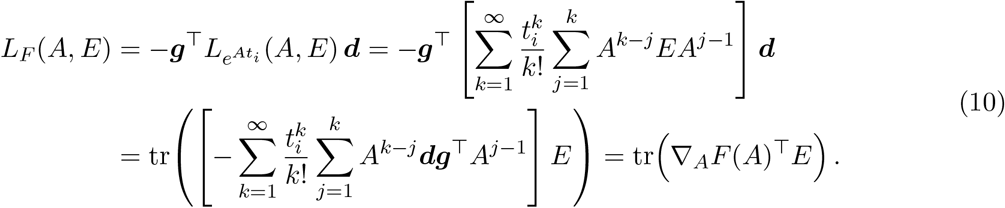

The first equality applies the chain rule to 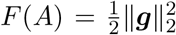. The second equality uses the Fréchet derivative 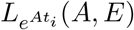 of the matrix exponential as given by Theorem 3.2 in [19]. The third equality follows from straightforward algebraic manipulations. Finally, by the definition of the matrix gradient as the unique matrix ∇_*A*_*F* (*A*) such that *L*_*F*_ (*A, E*) = tr ∇_*A*_*F* (*A*)^*T*^*E* for all *E*, we obtain

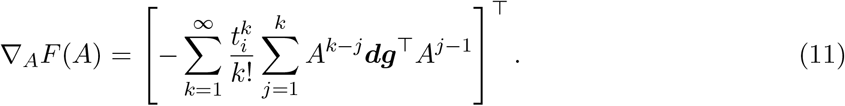

As shown in Eq. (11), ∇_*A*_*F* (*A*) is expressed as an infinite series. In practice, we can approximate this series using Algorithm 6.4 in [19] with the approximation error at the level of machine precision.

#### Time complexity

The Taylor apprxomiation of both (7) and (8) using (9) and (11) only involves a series of matrix-matrix multplications. Therefore, the overall time complexity is 𝒪 (*n*^3^).

### 3.5 Solve optimization (5) vid gradient descent

In the above sections, we introduce how we can efficiently compute the gradient 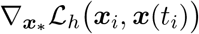and 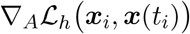. Then we optimize ***x***_*_ and *A* in Eq. (5) via gradient descent by updating 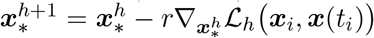 and 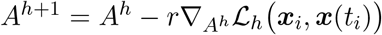, where *r* is the learning rate and *h* denotes the iteration index. These updates are repeated until convergence. Implementation details are in the supplementary materials.

#### Initialization

It is easy to verify that the optimization problem in (5) is non-convex. Therefore, initialization is crucial for gradient-based optimization. Throughout this paper, we initialize *A* = **0**_*n×n*_ in all experiments.

## 4 Finite–difference approximation vs. exact ODE solution

In this section, we provide a theoretical analysis of the gap between our exact estimator for the linear ODE in Eq. (2) and the estimator obtained from the first–order finite–difference approximation (1). For simplicity, we focus on the homogeneous system 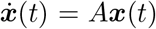,which is the same to Eq. (2) when ***x***_*_ = **0**.

The first–order finite–difference scheme approximates the time derivative by 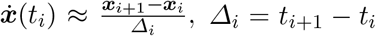,and thus replaces the ODE 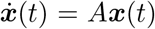 by the linear regression model 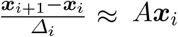. Instead of recovering the true matrix *A*, this approximation returns an estimate Â satisfying 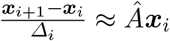. Concretely, Â is obtained as the minimizer of

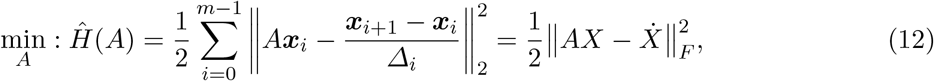

where *X* = [***x***_0_, …, ***x***_*m−*1_] collects the state vectors as columns and 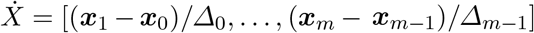.

In contrast, our methodology in Sections 3.1–3.4 directly estimates the ODE matrix *A* by fitting the exact solution of the linear system, 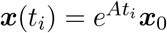, via

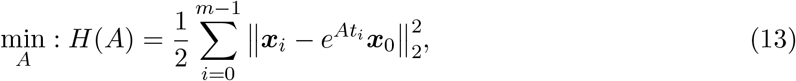

which is similar to our optimization (5) when *δ* is very large and ***x***_*_ = **0**.

The following theorem quantifies the theoretical gap between the optimal finite–difference solution of Eq. (12) and the exact-solution estimator in Eq. (13).

### Theorem 1

*Let* Â_*_ *denote the optimal finite–difference estimator obtained from Eq*. (12). *Then* Â_*_ *is in general a biased estimator of the true matrix A; more precisely*,

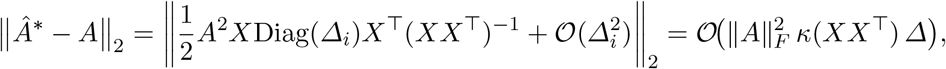

*where Δ* = max_*i*_ *Δ*_*i*_ *is the maximal time step and κ*(*XX*^*T*^) *is the (spectral) condition number*.

The proof of Theorem 1 is given in the supplementary materials. Theorem 1 shows that there is an inherent, non-vanishing bias between the finite–difference estimator and our exact estimator. The bias scales linearly with the maximum step size *Δ* and the condition number *κ*(*XX*^*T*^); in particular, larger time gaps and ill-conditioned *X* can significantly increase the discrepancy ‖ Â _*_ *− A* ‖ _2_.

Consequently, Theorem 1 provides theoretical support for the empirical observation that our exact method outperforms methods that rely on finite–difference approximations (such as SCODE, GRISLI, dynGENIE3, and PROB).

## 5 Experimental Results

In this section, we benchmark our method with existing methods on both simulation data and real-world single-cell RNA-seq data to demonstrate the outperformance of our method.

### Competing methods

To comprehensively benchmark performance, we compare our method against representative approaches from both dynamic and steady-state (non-temporal) gene regulation inference methods. On the dynamic side, we include ODE-based models: linear ODE-based methods (SCODE [8], GRISILI [9], and PROB [11]) and nonlinear/tree-based variants (dynGENIE3 [10]), as well as a causality-based model SWING [14] (reported as a top performer in the DREAM4 Network Inference Challenge for modeling expression dynamics [4]). On the steady-state side, we evaluate Bayesian network inference (IDA [9]), mutual information (MI [9]) methods, correlation (Cor [9]) method, ordinary least squares (LM mentioned as *lm* in [8]), lasso-regularized regression (Lasso mentioned as *msgps* in [8]), and regression-tree approaches JUMP3 [23] and GENIE3 [24] (the top performer in the DREAM5 Network Inference Challenge [2]).

### Evaluation metric

We evaluate each method using the Area Under the Receiver Operating Characteristic curve (AUROC) score, calculated by ranking TF-gene pairs by their predicted scores (e.g., confidence score or the absolute value of a coefficient) and comparing this ranking against the gold-standard TF-gene regulation interactions. We also compare the trajectories generated by *A*s learned by our method and SCODE, which are the only two methods that can recover *A*.

### Hyper-parameter tuning

Our model has two hyperparameters: the Huber threshold *δ* and the learning rate. We fix the learning rate at 10^−3^ for all experiments and select *δ* by cross-validation on each dataset. For each competing method, we perform a grid search with cross-validation to choose its hyperparameters. After selection, we fit each method with the chosen hyperparameters and report the resulting AUROC.

### 5.1 Benchmarking on simulation data

We first benchmark the competing methods on simulated data, where the ground-truth regulatory network between TF and their targets is known.

### Data simulation procedure

We simulate the time-course expression data {(**x**_*i*_, *t*_*i*_) ∈ ℝ ^*n*^ *×*ℝ :*i* = 1, …, *m*} from

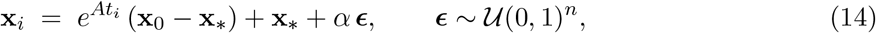

where ε is a noise vector with i.i.d. entries and *α* controls the noise level. Notably, when *α* = 0, Eq. (14) reduces to the noise-free solution of the linear ODE in Eq. (2).

Given the number of TFs *n*, time points *m*, and noise level *α >* 0, we can generate the simulated time-course expression data 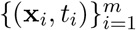 by following the procedure described below. First, we draw a random steady-state vector **x**_*_ ∈ [0, 8]^*n*^. Next, we sample a sparse interaction matrix *A* ∈ [−2, 2]^*n×n*^ with approximately 50% zero entries. To guarantee stability, we enforce strict diagonal dominance with negative diagonals: for each *i*, set *A*_*ii*_ *<* 0 such that |*A*_*ii*_| *>* Σ_*j*_ ≠ _*i*_ |*A*_*ij*_| + *κ* for a small *κ >* 0. By Gershgorin’s theorem [25], this construction makes *A* Hurwitz, and the system described in Eq. (14) converges. We then sample an initial condition **x**_0_ ∈ [0, 8]^*n*^ and *m* time points 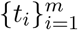with irregular intervals, where 0 *< t*_1_ *<* … *< t*_*m*_ *≤* 10. In the end, we substitute 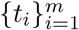, and 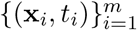 into Eq. (14) to yield the time-course data {(**x**_*i*_, *t*_*i*_)}^*m*^. The nonzero entries of *A* define the ground-truth regulatory network for benchmarking.

#### Simulation data

We use the above procedure to create two benchmark settings, **SD1** and **SD2**, that differ only in the number *m* of time points. In both settings, we fix *n* = 30 TFs and vary the noise level over *α* ∈ {0, 0.02, 0.04, 0.06, 0.08, 0.10}. For **SD1** we set *m* = 30, and for **SD2** we set *m* = 90. For each value of *α*, we generate 10 independent time-course datasets, yielding 60 datasets per setting.

#### Benchmarking results

We compare our method with dynamic gene regulation inference methods (ODE-based: SCODE, GRISILI, dynGENIE3, and PROB; Causality-based: SWING) on both **SD1** and **SD2**. Performance is assessed by the AUROC score for recovering the nonzero entries of the ground-truth interaction matrix *A*. Results are shown in Fig. 2(a–b). As illustrated, our method outperforms the competing methods for all settings in terms of AUROC scores. Furthermore, we observe that the performance of our method decreases as the noise level increases, which is as expected. However, we notice that the performance of our method on **SD2** (with more time points) is consistently better than its performance on **SD1** at each noise level, indicating that more data points can improve the performance of our method. By contrast, the competing struggle to recover *A* (AUROC near 0.5) and exhibit no clear dependence on the noise level.

**Fig. 2:**
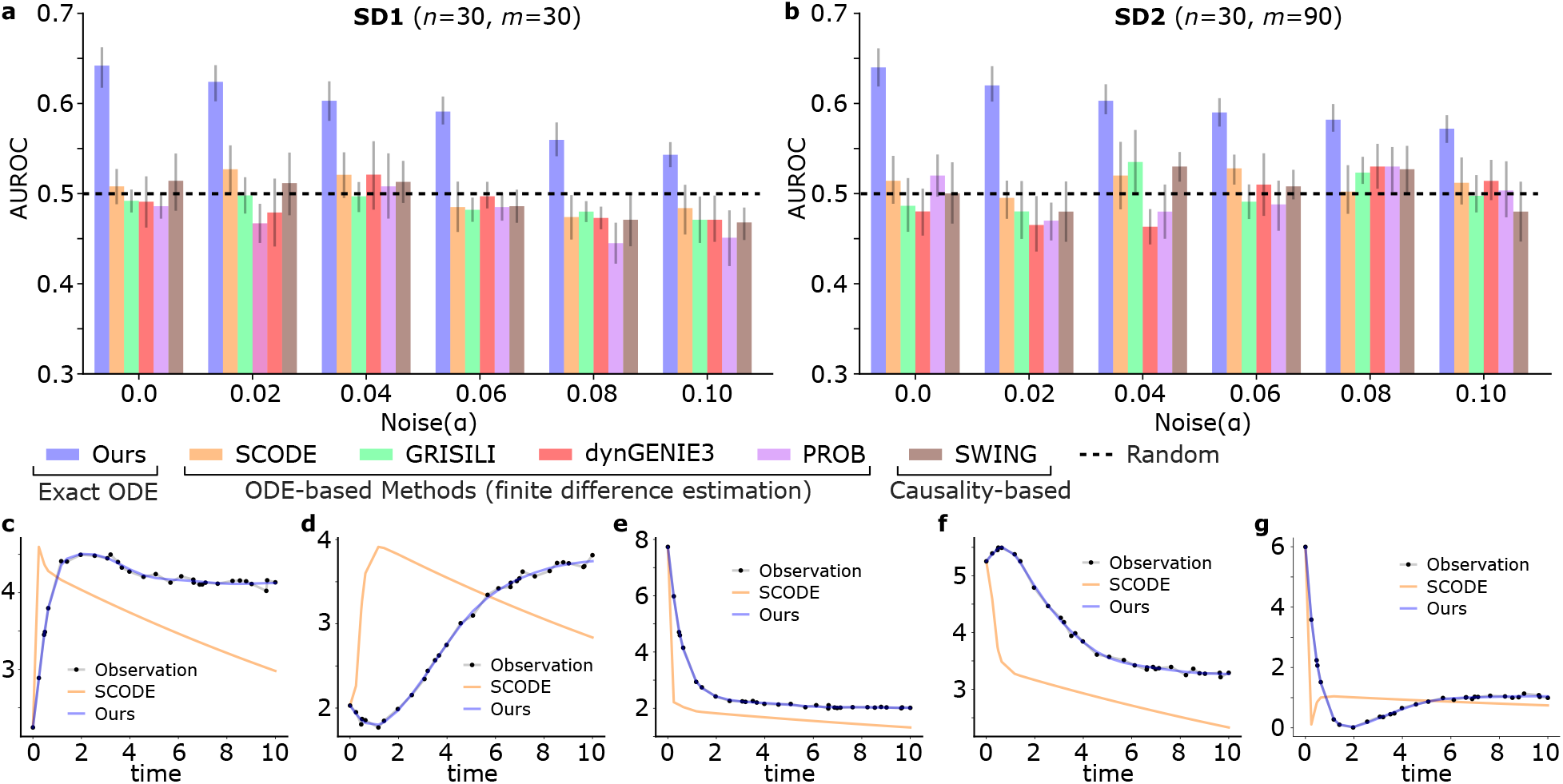
Benchmarking on simulation data. (a) AUROC on **SD1**. Bars show the mean AUROC and error bars denote the standard deviation across 10 independently simulated datasets at the same noise level *α*. (b) AUROC on **SD2**. (c–f) Representative time-course trajectories: observed data versus trajectories generated using *A* estimated by our method and by SCODE.

We further compare our method with SCODE on their ability to reproduce trajectories of the observed time-course expression in **SD1** when *α* = 0.04. Note that among the competing methods, only our method and SCODE are designed to estimate the numerical entries of the interaction matrix *A* in the linear ODE; other methods focus on ranking putative edges rather than recovering their values. After estimating *A* with each method, we generate the predicted trajectories from the learned models and compare them with the observed data. As shown in Fig. 2(c–g), trajectories produced using the *A* estimated by our method align more closely with the observations than those produced by SCODE, demonstrating superior dynamical fidelity.

### 5.2 Benchmarking on mouse MEFs–to–iN reprogramming data

#### Dataset

In this section, we evaluate competing methods on a scRNA-seq dataset profiling the direct reprogramming of mouse embryonic fibroblasts (MEFs) into induced neurons (iN) [26]. The dataset comprises 405 cells sampled at days 0, 2, 5, and 22. Cell-level pseudotime values were computed with Monocle [27] and provided by [26] as well. We benchmark all competing methods on inferring a TF regulatory network between 100 TFs as done by [8, 9]. The gold-standard regulatory network is extracted from the Transcription Factor Regulatory Network (TFRN) database [28, 29] for computing the AUROC.

#### Benchmarking results

We first run dynamic gene regulation inference methods using scRNA-seq data with the corresponding cell-level pseudotime. We further run steady-state gene regulation inference methods using only scRNA-seq data. The benchmarking results are shown in Fig. 3(a). Our method achieves the highest AUROC and outperforms all competing approaches by a clear margin.

**Fig. 3:**
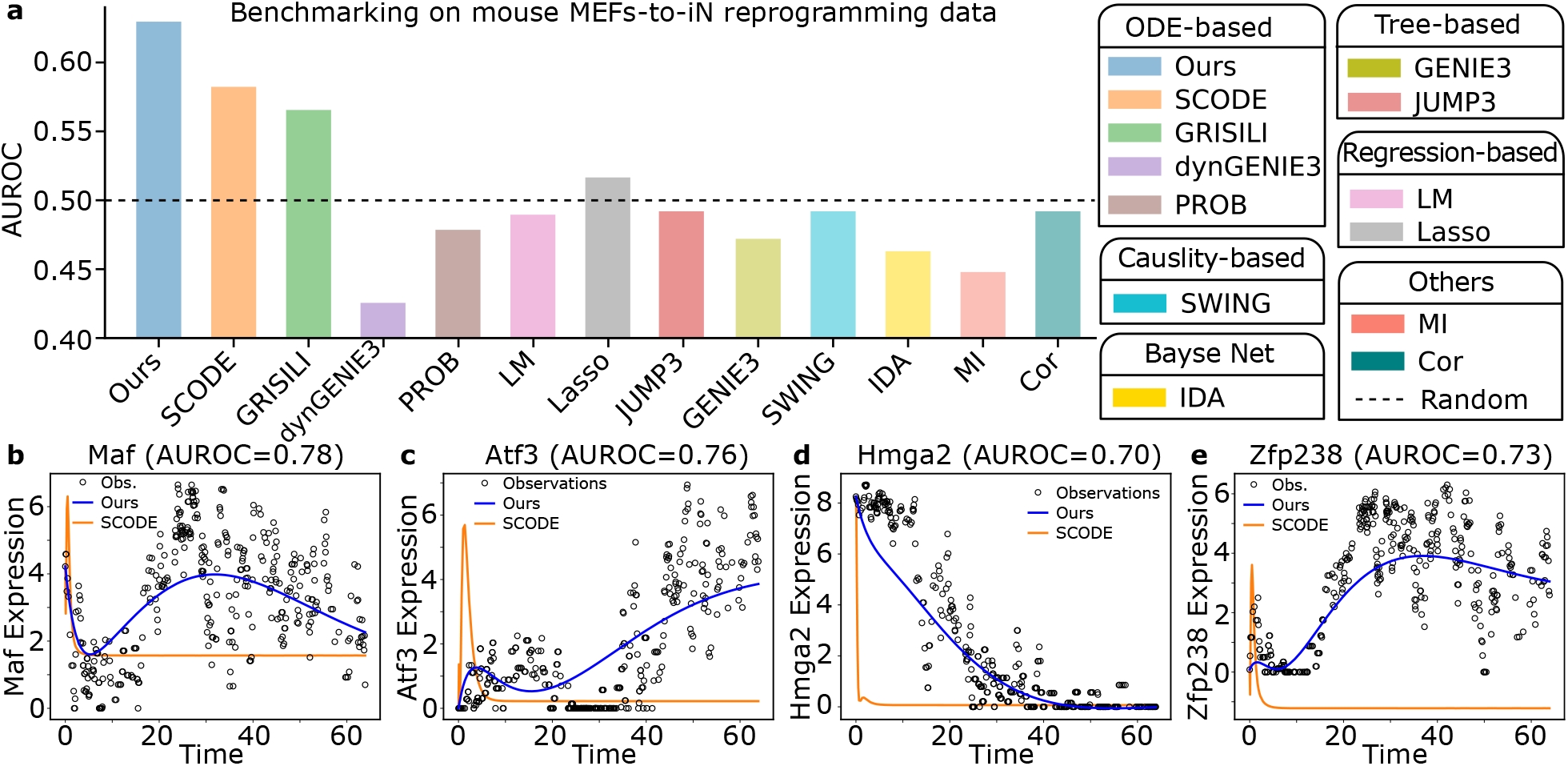
Benchmarking on mouse MEFs-to-iN reprogramming data. (a) AUROC comparision. (b–e) Trajectories generated using *A*s learned by our method and SCODE from observed expression data. Black circles are observed expression of the cells along pseudotime.

We further examine four representative TFs (Maf, Atf3, Hmga2, and Zfp238) for which our method attains high AUROC scores (0.78, 0.76, 0.70, and 0.73, respectively; Fig. 3b–e). For all four

TFs, the trajectories inferred by our model (blue) provide smooth fits that closely follow the main temporal patterns in the noisy observations, capturing the transient activation and subsequent decline of Maf, the gradual induction of Atf3, the monotonic down-regulation of Hmga2, and the sigmoidal activation of Zfp238. In contrast, SCODE (orange, the only competing method that can predict the trajectories) yields an early sharp spike followed by nearly constant expression levels, failing to reproduce the observed dynamics. These examples illustrate that our method not only recovers gold-standard regulators with higher AUROC but also reconstructs more biologically plausible TF expression trajectories from noisy scRNA-seq data.

### 5.3 Benchmarking on human ES cells to DE cells differentiation data

#### Dataset

In this section, we evaluate competing methods on a scRNA-seq dataset profiling differentiation of definitive endoderm (DE) cells from human embryonic stem (ES) cells [30]. The dataset comprises 758 cells sampled at 0, 12, 24, 36, 72, and 96 hours during the differentiation. Cell-level pseudotime is provided by [30] as well. We benchmark all competing methods on inferring a TF regulatory network between 100 TFs as done by [8, 9]. The gold-standard regulatory network is extracted from the TFRN database [28, 29] for computing the AUROC.

#### Benchmarking results

We benchmarked a panel of gene-regulation inference methods (as illustrated in the legend of Fig. 4) on the human ES cells to DE cells differentiation data scRNA-seq dataset. Figure 4(a) summarizes AUROC scores against the gold-standard regulatory network, with the dashed line indicating random performance (AUROC = 0.5). Our method achieves the highest AUROC and outperforms all competing approaches by a clear margin.

**Fig. 4:**
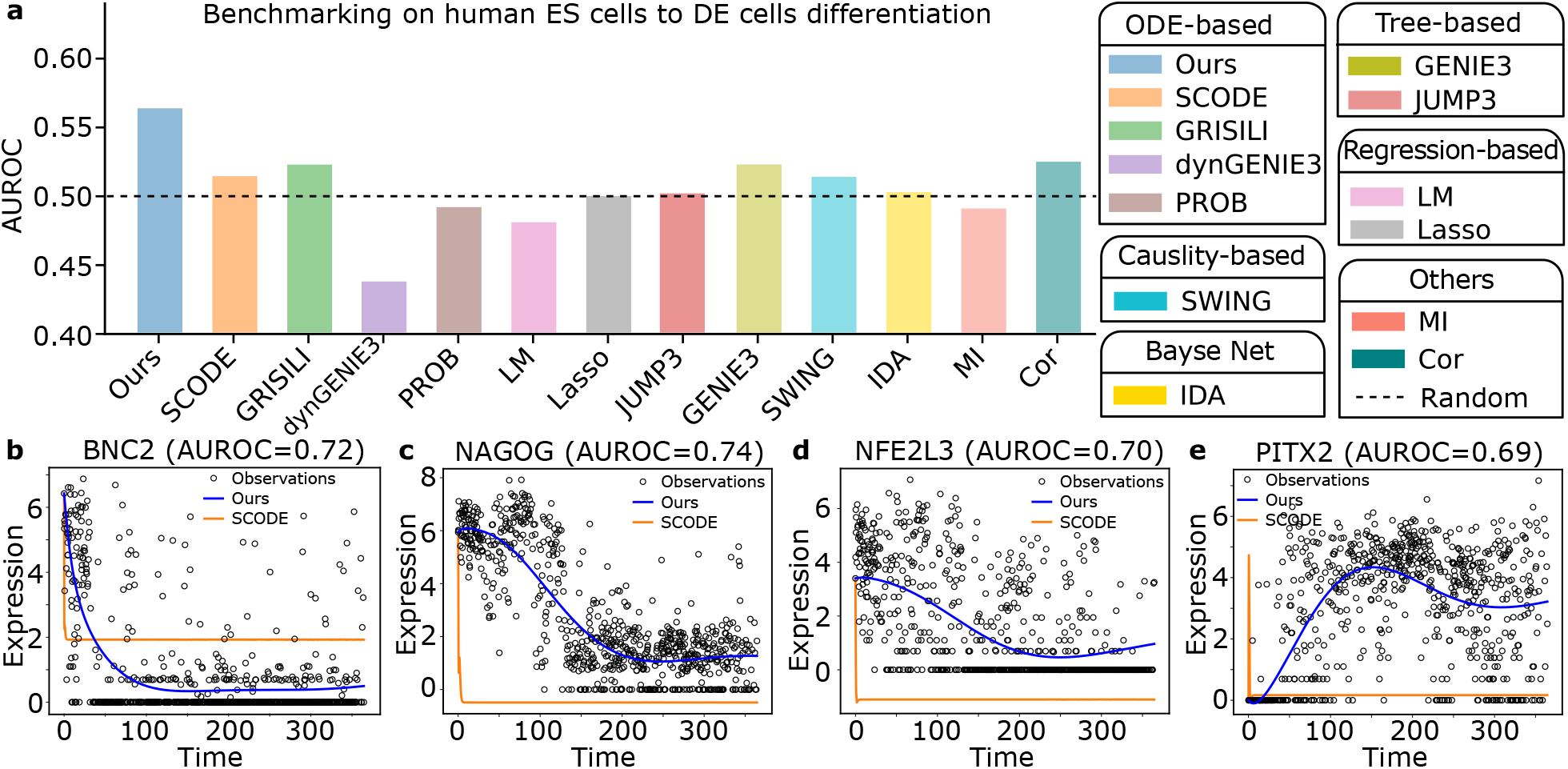
Benchmarking on human ES cells to DE cells differentiation data. (a) AUROC comparision. (b–e) Trajectories generated using *A*s learned by our method and SCODE from observed expression data. Black circles are observed expression of the cells along pseudotime.

We further illustrate four representative TFs (BNC2, NAGOG, NFE2L3, and PITX2), for which our method achieves high AUROC scores (0.72, 0.74, 0.70, and 0.69, respectively; Fig. 4b–e). Similar to what we observed in Fig. 3, trajectories inferred by our model fits better to the noisy single-cell observations than SCODE.

### 5.4 Benchmarking on mouse ES cells to PrE cells differentiation data

#### Dataset

In this section, we evaluate competing methods on a scRNA-seq dataset profiling differentiation of primitive endoderm (PrE) cells from mouse embryonic stem (ES) cells [8]. The dataset comprises 456 cells sampled at 0, 12, 24, 48, and 72 hours during the differentiation. Cell-level pseudotime is provided by [8] as well. We benchmark all competing methods on inferring a TF regulatory network between 100 TFs as done by [8]. The gold-standard regulatory network is extracted from the TFRN database [28, 29] for computing the AUROC.

#### Benchmarking results

We benchmarked a panel of gene-regulation inference methods (as illustrated in the legend of Fig. 5) on the mouse ES cells to PrE cells differentiation data scRNA-seq dataset. Figure 5(a) summarizes AUROC scores against the gold-standard regulatory network, with the dashed line indicating random performance (AUROC = 0.5). Our method achieves the highest AUROC and outperforms all competing approaches by a clear margin.

**Fig. 5:**
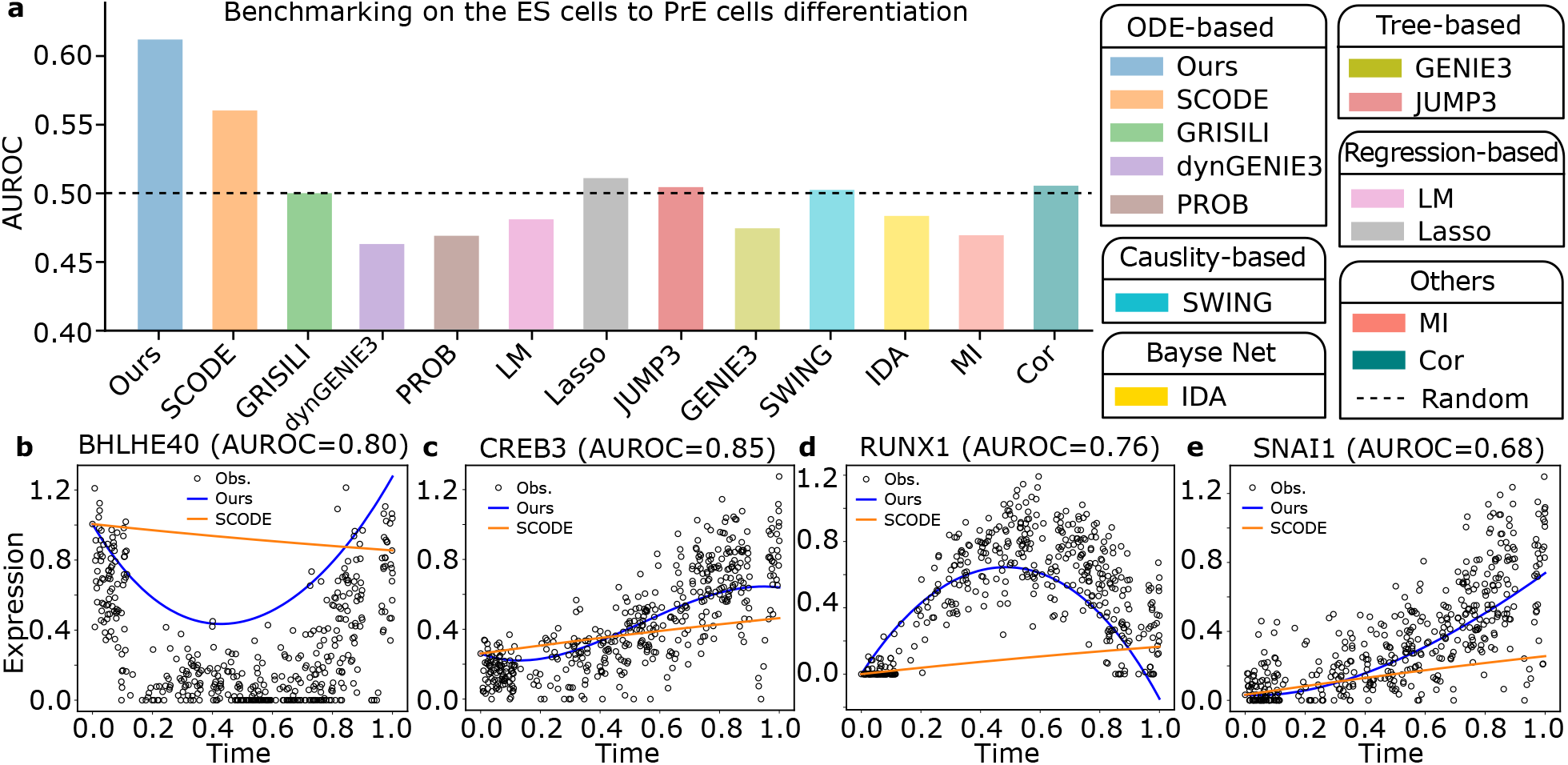
Benchmarking on mouse ES cells to PrE cells differentiation data. (a) AUROC comparision. (b–e) Trajectories generated using *A*s learned by our method and SCODE from observed expression data. Black circles are observed expression of the cells along pseudotime.

We further illustrate four representative TFs (BHLHE40, CREB3, RUNX1, and SNAI1), for which our method achieves high AUROC scores (0.80, 0.85, 0.76, and 0.68, respectively; Fig. 5b–e). Similar to what we observed in Figs. 3 and 4, trajectories inferred by our model fit better to the noisy single-cell observations than SCODE.

### 5.5 Runtime of our method

On simulation data, our method takes about 1 minute; on the scRNA-seq datasets in Sections 5.2 and 5.4, the runtime is around 6 minutes; and on the scRNA-seq dataset in Section 5.3, the runtime is around 10 minutes.

## 6 Conclusions

In this paper, we introduce an efficient method to directly solve linear ODE models without relying on finite-difference approximations. We apply this method to infer gene regulatory networks from time-course gene expression data and demonstrate that it outperforms existing approaches in both gene regulation inference and expression dynamics modeling.

## Acknowledgment

This research was supported by National Institutes of Health [R35GM147241 to Y.W.]. This research was also supported in part by the Division of Intramural Research of the National Library of Medicine (NLM), National Institutes of Health (NIH). The contributions of the NIH author are considered Works of the United States Government. The findings and conclusions presented in this paper are those of the authors and do not necessarily reflect the views of the NIH or the U.S. Department of Health and Human Services.

## Footnotes

5 If ‖*A*‖_2_ is large, we use *scaling and squaring* (see [19] and Chapter 11.3.1 [21]): first we find *s* such that ‖*A/*2^*s*^‖*≤* 1, then apply degree-18 Taylor approximation on exp(*A*) = exp(*A/*2^s^)^s^.

## References

[1] L. Garcia-Alonso et al. “Benchmark and integration of resources for the estimation of human transcription factor activities”. In: Genome Res 29.8 (Aug. 2019), pp. 1363–1375.

[2] Daniel Marbach et al. “Wisdom of crowds for robust gene network inference”. In: Nature Methods 9.8 (2012), pp. 796–804.

[3] André Fujita et al. “Modeling gene expression regulatory networks with the sparse vector autoregressive model”. en. In: BMC Syst. Biol. 1.1 (Aug. 2007), p. 39.

[4] Jonathan Lu et al. “Causal network inference from gene transcriptional time-series response to glucocorticoids”. en. In: PLoS Comput. Biol. 17.1 (Jan. 2021), e1008223.

[5] Shun Yao, Shinjae Yoo, and Dantong Yu. “Prior knowledge driven Granger causality analysis on gene regulatory network discovery”. en. In: BMC Bioinformatics 16.1 (Aug. 2015), p. 273.

[6] Liang Chen et al. “Inferring gene regulatory networks from time-series scRNA-seq data via GRANGER causal recurrent autoencoders”. en. In: Brief. Bioinform. 26.2 (Mar. 2025).

[7] Atul Deshpande et al. “Network inference with Granger causality ensembles on single-cell transcriptomics”. en. In: Cell Rep. 38.6 (Feb. 2022), p. 110333.

[8] Hirotaka Matsumoto et al. “SCODE: an efficient regulatory network inference algorithm from single-cell RNA-Seq during differentiation”. en. In: Bioinformatics 33.15 (Aug. 2017), pp. 2314–2321.

[9] Pierre-Cyril Aubin-Frankowski and Jean-Philippe Vert. “Gene regulation inference from single-cell RNA-seq data with linear differential equations and velocity inference”. en. In: Bioinformatics 36.18 (Sept. 2020), pp. 4774–4780.

[10] V ân Anh Huynh-Thu and Pierre Geurts. “dynGENIE3: dynamical GENIE3 for the inference of gene networks from time series expression data”. en. In: Sci. Rep. 8.1 (Feb. 2018), p. 3384.

[11] Xiaoqiang Sun, Ji Zhang, and Qing Nie. “Inferring latent temporal progression and regulatory networks from cross-sectional transcriptomic data of cancer samples”. en. In: PLoS Comput. Biol. 17.3 (Mar. 2021), e1008379.

[12] Atte Aalto et al. “Gene regulatory network inference from sparsely sampled noisy data”. en. In: Nat. Commun. 11.1 (July 2020), p. 3493.

[13] Andrea Ocone et al. “Reconstructing gene regulatory dynamics from high-dimensional single-cell snapshot data”. en. In: Bioinformatics 31.12 (June 2015), pp. i89–96.

[14] Justin D Finkle, Jia J Wu, and Neda Bagheri. “Windowed Granger causal inference strategy improves discovery of gene regulatory networks”. en. In: Proc. Natl. Acad. Sci. U. S. A. 115.9 (Feb. 2018), pp. 2252–2257.

[15] Hirotaka Matsumoto and Hisanori Kiryu. “SCOUP: a probabilistic model based on the Ornstein-Uhlenbeck process to analyze single-cell expression data during differentiation”. en. In: BMC Bioinformatics 17.1 (June 2016), p. 232.

[16] Mukesh Bansal, Giusy Della Gatta, and Diego di Bernardo. “Inference of gene regulatory networks and compound mode of action from time course gene expression profiles”. en. In: Bioinformatics 22.7 (Apr. 2006), pp. 815–822.

[17] Intekhab Hossain et al. “Biologically informed NeuralODEs for genome-wide regulatory dynamics”. en. In: Genome Biol. 25.1 (May 2024), p. 127.

[18] Maggie Beheler-Amass et al. “Dynamic gene regulatory network Inference with interpretable, biophysically-motivated neural ODEs”. Sept. 2025.

[19] Awad H Al-Mohy and Nicholas J Higham. “Computing the Fr échet Derivative of the Matrix Exponential, with an application to Condition Number Estimation”. In: SIAM Journal on Matrix Analysis and Applications 30.4 (2009), pp. 1639–1657.

[20] Michael S Paterson and Larry J Stockmeyer. “On the Number of Nonscalar Multiplications Necessary to Evaluate Polynomials”. In: SIAM Journal on Computing 2.1 (1973), pp. 60–66.

[21] Gene H Golub and Charles F Van Loan. Matrix Computations. JHU press, 2013.

[22] Philipp Bader, Sergio Blanes, and Fernando Casas. “Computing the Matrix Exponential with an Otimized Taylor Polynomial Approximation”. In: Mathematics 7.12 (2019), p. 1174.

[23] V â n Anh Huynh-Thu and Guido Sanguinetti. “Combining tree-based and dynamical systems for the inference of gene regulatory networks”. en. In: Bioinformatics 31.10 (May 2015), pp. 1614–1622.

[24] V. A. Huynh-Thu et al. “Inferring regulatory networks from expression data using tree-based methods”. In:PLoS One 5.9 (Sept. 2010).

[25] Roger A Horn and Charles R Johnson. Matrix Analysis. en. 2nd ed. Cambridge, England: Cambridge University Press, Oct. 2012.

[26] Barbara Treutlein et al. “Dissecting direct reprogramming from fibroblast to neuron using single-cell RNA-seq”. en. In: Nature 534.7607 (June 2016), pp. 391–395.

[27] Cole Trapnell et al. “The dynamics and regulators of cell fate decisions are revealed by pseudotemporal ordering of single cells”. en. In: Nat. Biotechnol. 32.4 (Apr. 2014), pp. 381–386.

[28] Shane Neph et al. “Circuitry and dynamics of human transcription factor regulatory networks”. en. In: Cell 150.6 (Sept. 2012), pp. 1274–1286.

[29] Andrew B Stergachis et al. “Conservation of trans-acting circuitry during mammalian regulatory evolution”. en. In: Nature 515.7527 (Nov. 2014), pp. 365–370.

[30] Li-Fang Chu et al. “Single-cell RNA-seq reveals novel regulators of human embryonic stem cell differentiation to definitive endoderm”. en. In: Genome Biol. 17.1 (Dec. 2016).

[31] Nicholas J Higham. Functions of Matrices: Theory and Computation. SIAM, 2008.

